# The Okur-Chung Neurodevelopmental Syndrome (OCNDS) mutation CK2^K198R^ leads to a rewiring of kinase specificity

**DOI:** 10.1101/2021.04.05.438522

**Authors:** Danielle M Caefer, Nhat Q Phan, Jennifer C Liddle, Jeremy L Balsbaugh, Joseph P O’Shea, Anastasios V Tzingounis, Daniel Schwartz

## Abstract

Okur-Chung Neurodevelopmental Syndrome (OCNDS) is caused by heterozygous mutations to the CSNK2A1 gene, which encodes the alpha subunit of casein kinase II (CK2). The most frequently occurring mutation is lysine 198 to arginine (K198R). To investigate the impact of this mutation, we first generated a high-resolution phosphorylation motif of CK2^WT^, including the first characterization of specificity for tyrosine phosphorylation activity. A second high resolution motif representing CK2^K198R^ substrate specificity was also generated. Here we report for the first time the impact of the OCNDS associated CK2^K198R^ mutation. Contrary to prior speculation, the mutation does not result in a loss of function, but rather shifts the substrate specificity of the kinase. Broadly speaking the mutation leads to 1) a decreased preference for acidic residues in the +1 position, 2) a decreased preference for threonine phosphorylation, 3) an increased preference for tyrosine phosphorylation, and 4) an alteration of the tyrosine phosphorylation specificity motif. To further investigate the result of this mutation we have developed a probability-based scoring method, allowing us to predict shifts in phosphorylation in the K198R mutant relative to the wild type kinase. As an initial step we have applied the methodology to the set of axonally localized ion channels in an effort to uncover potential alterations of the phosphoproteome associated with the OCNDS disease condition.

## Introduction

Okur-Chung Neurodevelopmental Syndrome (OCNDS) is broadly characterized by delayed psychomotor development and intellectual disability.^1^ The disorder has been linked to various heterozygous mutations on the CSNK2A1 gene, which encodes the alpha subunit of casein kinase II (CK2), a well-characterized and conserved serine/threonine protein kinase.^2,3^ Of the mutations associated with OCNDS, the mutation of lysine-198 to arginine (K198R) has been most frequently observed and has been speculated to result in a loss of kinase function.^4^ Interestingly, the K198R mutation is located in the activation segment of CK2, a highly conserved region important for kinase activity and substrate recognition.^5^ The substrate specificity of wild type CK2 is well-characterized with a clear preference for acidic residues in the +1 and +3 residues relative to the phosphoacceptor.^6^ Lysine-198 is thought to interact with these acidic amino acids of substrate proteins downstream of the phosphoacceptor residue^6^ and our group previously posited that the residue would specifically be involved in the specificity of the +1 position.^7^ To determine the impact of the CK2^K198R^ mutation we implemented the ProPeL method to generate a high-resolution motif of CK2^WT^ and CK2^K198R^.^8^ Additionally, we utilized a probabilistic strategy to determine the differential likelihood of modification for serine, threonine, and tyrosine residues on axonally localized ion channels under wild type and mutant kinase conditions.

## Methods

### Plasmids

For bacterial expression, a plasmid containing the human CSNK2A1 gene in an Invitrogen Gateway donor vector (pDONR223) was provided by The Broad Institute and was transferred to the pDEST17 backbone following the standard Gateway Protocol (Life Technologies). Generation of the CK2^K198R^ mutant was done using the Q5® Site-Directed Mutagenesis Kit (New England BioLabs).

### ProPeL Experiments for Motif Determination

ProPeL experiments were carried out as previously described^8^ with the following conditions for *in vivo* proteome phosphorylation: all CK2 constructs were expressed in *Escherichia coli* OverExpress C43(DE3) cells (Lucigen) by IPTG induction. Optimal expression conditions were determined to be mid-log induction followed by expression for 24h at 37°C in TB media (data not shown).

### Untargeted Protein Identification via Tandem Mass Spectrometry

Peptide samples were subjected to mass analysis using a Thermo Scientific Ultimate 3000 RSLCnano ultra-high performance liquid chromatography (UPLC) system coupled to a high-resolution Thermo Scientific Q Exactive HF mass spectrometer. An aliquot of each peptide preparation in Solvent A (0.1% formic acid in H_2_O) was injected onto a Waters nanoEase m/z Peptide BEH C18 analytical column (130A, 1.7 μm, 75 μm × 250 mm) and separated by reverse-phase UPLC using a gradient of 4-30% Solvent B (0.1% formic acid in acetonitrile) over a 100-min gradient at 300 nL/min flow. Peptides were eluted directly into the Q Exactive HF using positive mode nanoflow electrospray ionization and 1.5 kV capillary voltage. MS scan acquisition parameters included 60,000 resolution, 1e6 AGC target, maximum ion time of 60 ms, and a 300 to 1800 m/z mass range. Data-dependent MS/MS scan acquisition parameters included 15,000 resolution, 1e5 AGC target, maximum ion time of 40 ms, loop count of 15, isolation window of 2.0 m/z, dynamic exclusion window of 30 s, normalized collision energy of 27, and charge exclusion “on” for all unassigned, +1, and >+8 charged species.

Peptides were identified using MaxQuant (v1.6.10.43) and the embedded Andromeda search engine.^9^ The raw data was searched against three databases: an in-house-generated protein database consisting of 6xHis-tagged CK2 wildtype and mutant sequences, the complete UniProt *E. coli* reference proteome (identifier UP0000068040, accessed 11Sept2020), and the MaxQuant contaminants database. Variable modifications were oxidation of Met, acetylation of protein N-termini, deamidation of Asn/Gln, and for enriched samples, phosphorylation of Ser/Thr/Tyr. Carbamidomethylation of Cys was set as a fixed modification. Protease specificity was set to trypsin, allowing a maximum of 2 missed cleavages. LFQ quantification was enabled. All results were filtered to a 1% false discovery rate at the peptide spectrum match and protein levels; all other parameters were kept at default values. MaxQuant-derived output was further analyzed in accordance with the ProPeL method.^10^

### Probability-based prediction of CK2^WT^ and CK2^K198R^ phosphorylation sites

Given that the ProPeL methodology for kinase motif determination is carried out in the context of a closed system (the *E. coli* expressed proteome) the scoring of phosphorylated and unphosphorylated serine, threonine, and tyrosine residues in *E. coli* could be used to convert motif scores to probabilities of modification. Specifically, phosphorylated 15mers (for either the wild type or K198R data set) were mapped back onto the UniProt reference K12 *E. coli* proteome to generate a list of expressed proteins phosphorylated by CK2 in *E. coli*. These proteins were then subjected to a full trypsin digestion *in silico* and resultant peptides with a length up to 50 residues were retained to generate a list of all potential phosphorylation sites that could have been hypothetically observed in our experiments. Seven residues upstream and downstream from each serine, threonine, and tyrosine residue in these peptides was extracted to create a complete list of serine, threonine, and tyrosine-centered 15mers. Each 15mer was labeled as either positive or negative depending on whether it was observed to be phosphorylated in our ProPeL experiments, resulting in 1350 positive (phosphorylated) and 26331 negative (not observed to be phosphorylated) 15mers in the CK2^WT^ data set, and 1024 positive and 20083 negative 15mers in the CK2^K198R^ data set. Every peptide was scored using the appropriate wild type or K198R serine, threonine, or tyrosine-centered position weight matrix derived from the log-odds binomial probabilities observed in the corresponding serine, threonine, or tyrosine-centered pLogo (as we have shown previously^11^). Note, the value for the central phosphorylated residue was derived from pLogo containing phosphorylated serine, threonine, and tyrosine-centered 15mers relative to presumed unphosphorylated serine, threonine, and tyrosine-centered 15mers (i.e., the pLogo with the unfixed central residue, see **Figure 1**). The complete set of scored 15mers was sorted in descending order and was used as a reference table to determine probabilities of modification based on score - specifically by taking the number of positive peptides above a score threshold divided by the total number of positive and negative peptides above a score threshold. Complete reference tables for CK2^WT^ and CK2^K198R^ are available in **Table S5** and **Table S6**. This methodology was utilized to score and assign probabilities of modification to all serine, threonine, and tyrosine residues in human axonally localized ion channels (see **Table 1** and **Table S7**).

**Figure 1.**
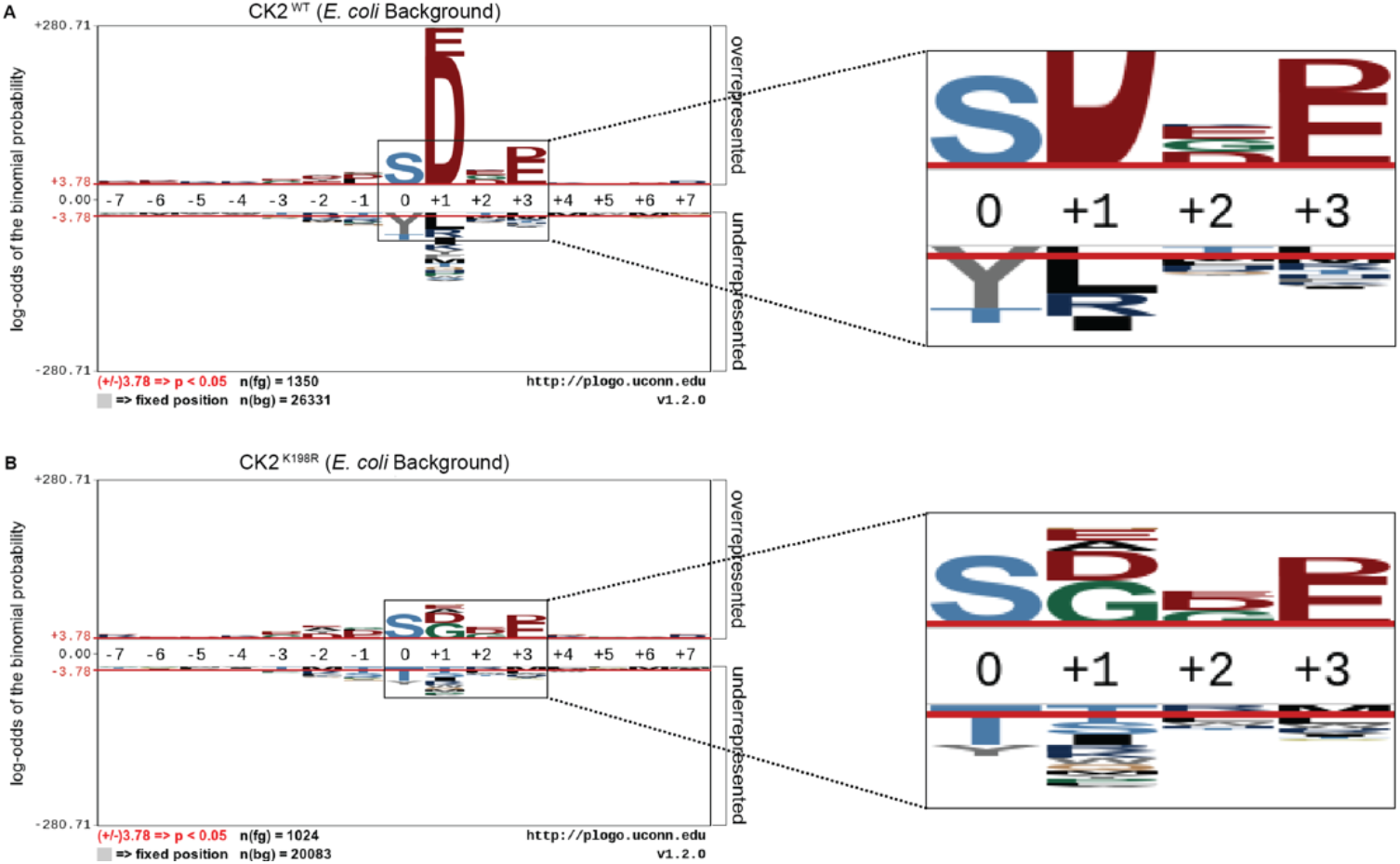
CK2^K198R^ mutation shifts overall specificity motif. Substrate specificity of CK2^WT^ **(A)** and CK2^K198R^ **(B)**. The CK2^K198R^ mutation results in a decreased preference for acidic (D/E) residues in the +1 position as well as a reordering of phosphoacceptor preference. In CK2^K198R^, a decreased preference for threonine is paired with an increase in preference for tyrosine.

**Table 1.**
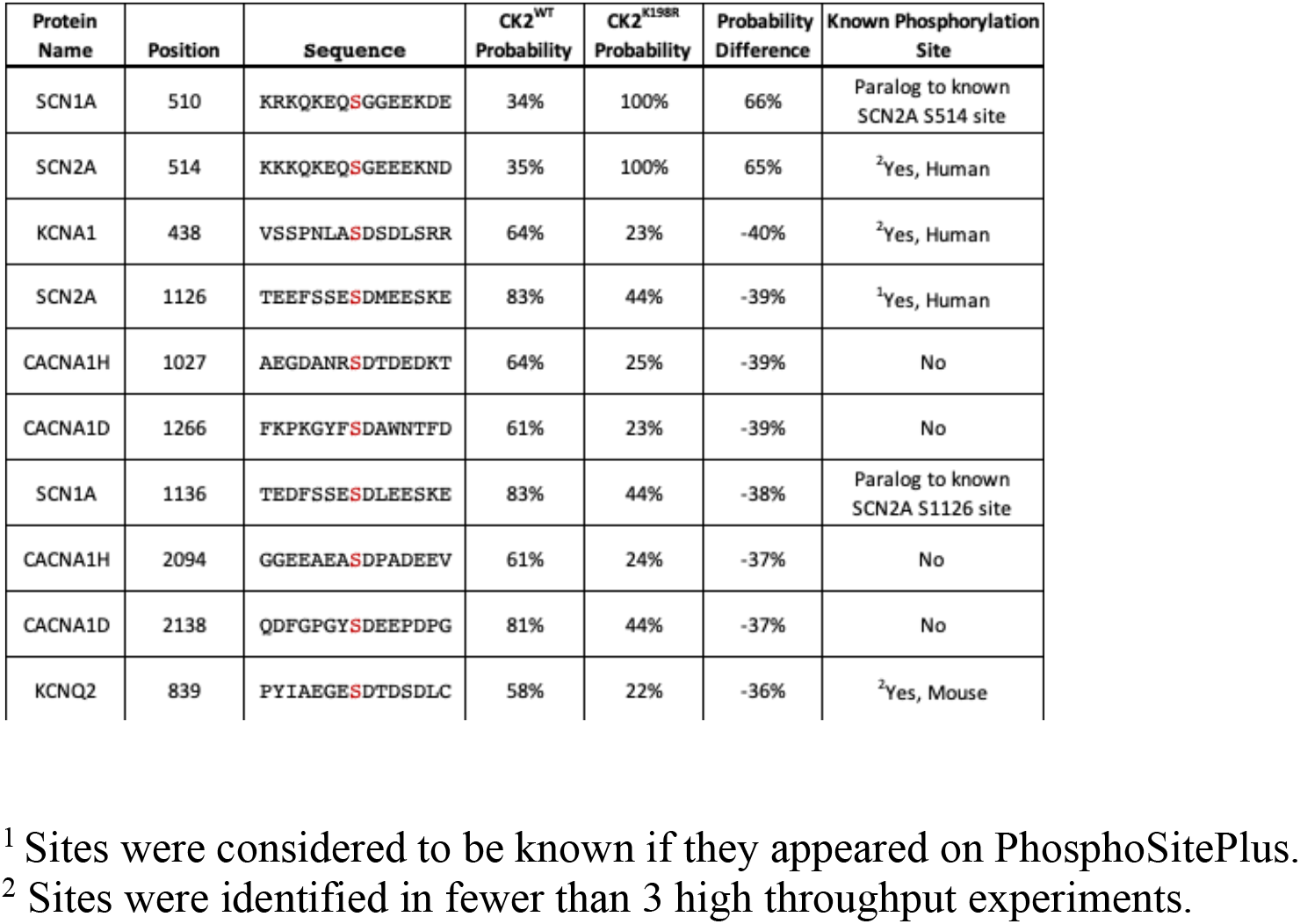
Top ten phosphorylation sites predicted to be most differentially phosphorylated by CK2^WT^ and CK2^K198R^ in axonally localized human ion channels.

## Results

### CK2^K198R^ is an active kinase

To obtain a higher resolution motif than we previously published in 2012^10^ for CK2^WT^, several new ProPeL experiments were carried out utilizing more sensitive instrumentation that allowed confident identification of low abundance phosphorylated peptides that escaped identification previously. Experiments with CK2^WT^ identified 3894 tryptic phosphopeptides (**Table S1**) with 2845 phosphopeptides classified as high confidence (i.e., containing at least one phosphorylation site with greater than 0.9 probability of modification determined by MaxQuant/Andromeda). Parallel ProPeL experiments with CK2^K198R^ identified 3322 (**Table S2**) tryptic phosphopeptides with 2439 phosphopeptides classified as high confidence. Each high confidence phosphorylation site was mapped back to the UniProt reference K12 *E. coli* proteome and extended to create lists of unique 15mers as previously described.^8^ Finally, 312 phosphorylation sites detected by our group and others as endogenous to *E. coli* ^7,10,12,13^ were removed from each list prior to motif analysis. This process yielded a data set comprised of 1350 unique phosphorylation sites (818 pS, 422 pT, 110 pY) for CK2^WT^ (**Table S3**) and 1024 unique phosphorylation sites (619 pS, 251 pT, 154 pY) for CK2^K198R^ (**Table S4**). Taken together these results indicate that CK2^K198R^ is an active kinase able to phosphorylate substrates under *in vivo* conditions. Furthermore, given that the CK2^WT^ and CK2^K198R^ ProPeL experiments were carried out under identical conditions, both the number of phosphorylation sites identified and comparable phosphopeptide intensities observed (data not shown) could be used as a proxy for kinase activity, thus suggesting that the OCNDS CK2^K198R^ mutant possesses a similar level of kinase activity to wild type CK2. This observed activity of CK2^K198R^ stands in contrast to previous speculation that the mutation may result in a loss of activity.^4^

### CK2^K198R^ exhibits an altered substrate specificity

The generation of pLogos^14^ provides a visual representation of the relative statistical significance of amino acids (represented by single letter abbreviations) at positions within a 15-residue window of a central phosphoacceptor residue. Amino acid preferences at each position are stacked and sized according to significance relative to the background data set. The most significant residues are displayed with the largest size and are positioned closest to the x-axis. Background data sets representing unphosphorylated residues were created independently for CK2^WT^ and CK2^K198R^. Each background was generated through the *in silico* tryptic digestion of proteins identified by the presence of at least one phosphorylated peptide in either the CK2^WT^ or CK2^K198R^ experiments. Background lists included serine, threonine, and tyrosine sites in proteins known to be expressed that could potentially have been observed via MS/MS, but were never detected in a phosphorylated state. These sites were subsequently extended 7 amino acids upstream and downstream of each phosphorylatable residue to generate two unique background lists for use in pLogo generation.

The overall pLogo generated with the current CK2^WT^ data set (**Figure 1A**) recapitulated our previously published CK2^WT^ pLogo^10^; namely a strong preference for acidic residues (D/E) at the +1 and +3 positions with the +1 position being the most statistically significant position in the pLogo overall. Consistent with our prior hypothesis that the K198R mutation would impact the specificity of the +1 position, the pLogo generated for CK2^K198R^ indicated a massively decreased preference for acidic (D/E) residues in the +1 position, as well as an increased preference for both glycine and alanine, with glycine superseding aspartate to become the most statistically significant residue at the +1 position (**Figure 1B**). Interestingly, the reduction in significance was limited to the +1 position, as the preference for acidic residues at the +3 position was maintained at nearly identical levels (compare **Figure 1A, 1B**).

Further comparison of the CK2^WT^ and CK2^K198R^ derived pLogos indicated an unexpected reordering of phosphoacceptor preference. Though both mutant and wild type CK2 strongly favored serine phosphorylation over threonine or tyrosine phosphorylation, CK2^WT^ strongly disfavored tyrosine as a phosphoacceptor and was relatively neutral towards threonine as a phosphoacceptor (**Figure 1A**), while CK2^K198R^ strongly disfavored threonine as a phosphoacceptor and was relatively neutral towards tyrosine as a phosphoacceptor. It is important to note that although CK2^K198R^ did not display a preference for tyrosine phosphorylation (i.e., it was not phosphorylated by CK2^K198R^ at a level greater than its relative frequency among phosphorylatable residues), its shift is significant as it represents a near doubling of the proportion of phosphorylated tyrosine residues in the K198R data set (15%) relative to wild type (8.1%).

### CK2^K198R^ exhibits altered specificity at the phosphoacceptor level

We next investigated whether the decreased preference for acidic residues at the +1 position observed in the overall CK2^K198R^ pLogo was consistent among each individual phosphoacceptor. To investigate this question, we fixed each central residue (serine, threonine, and tyrosine) to generate phosphoacceptor specific pLogos as shown in **Figure 2**. As was observed in the overall pLogo, the serine and threonine centered CK2^WT^ pLogos indicated a preference for acidic residues at the +1 and +3 positions (**Figure 2A, 2B**). The preference for acidic residues was only significant at the +1 position of the tyrosine centered pLogo (**Figure 2C**). In the CK2^K198R^ serine, threonine, and tyrosine centered pLogos (**Figure 2D-F**), the decreased preference for acidic residues at the +1 position was preserved. Interestingly however, the tyrosine centered pLogo suggested a modest increased preference for acidic residues at the −1 and −2 positions (**Figure 2F**). Importantly, the pLogos shown in **Figures 2C** and **2F** represent the first-ever documentation of a tyrosine motif specificity for CK2.

**Figure 2.**
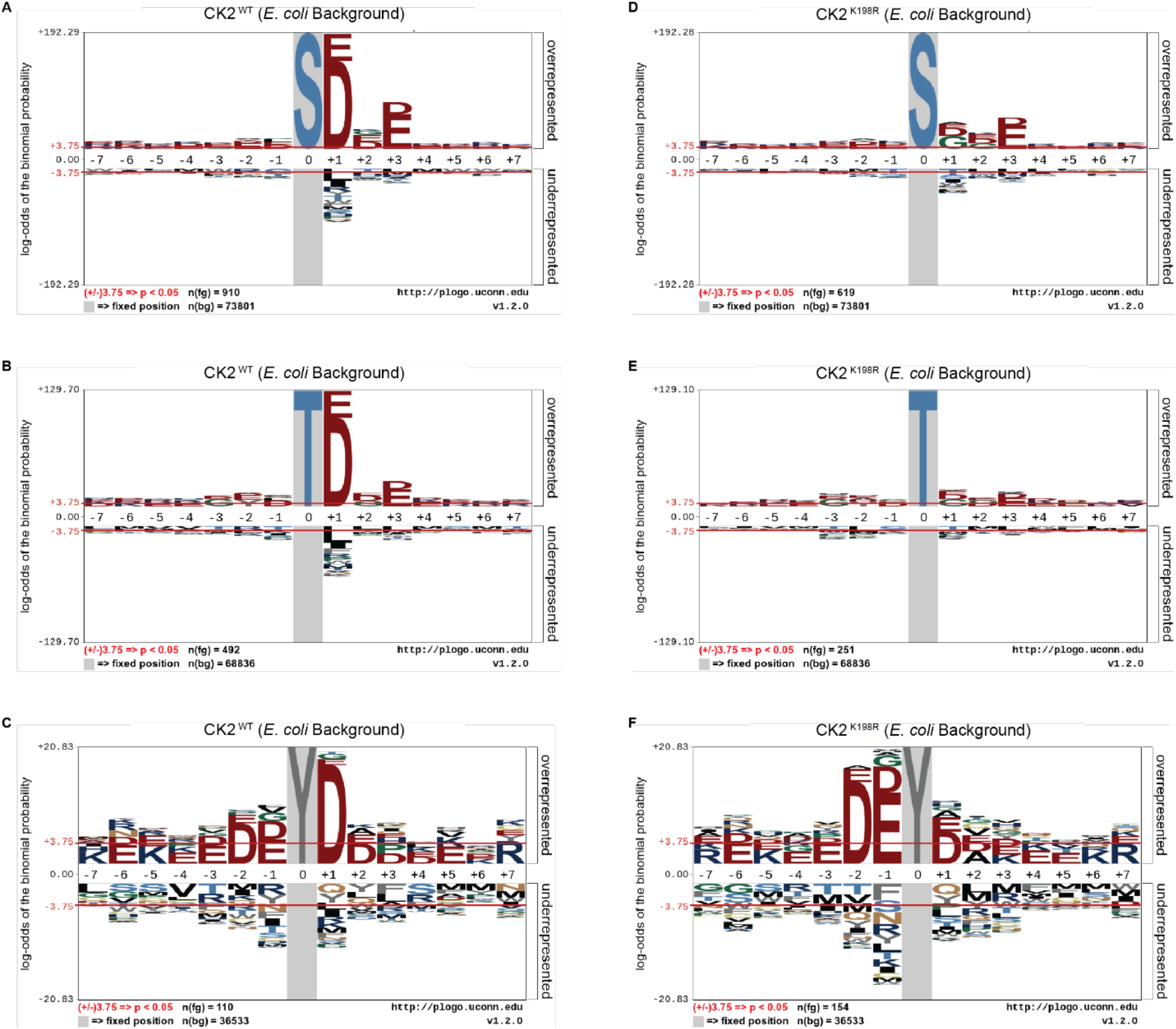
CK2^K198R^ exhibits altered +*1* substrate specificity. Substrate specificity of CK2^WT^ (**A-C**) and CK2^K198R^(**D-F**). The CK2^K198R^ mutation results in a decreased preference for acidic (D/E) residues in the +1 position for S and T centered phosphorylation sites. For Y centered phosphorylation sites, the preference for acidic residues is shifted to the −2 and −1 positions.

### CK2^K198R^ mutation substantially alters the landscape of CK2 phosphorylation at the site level

The ProPeL methodology, carried out in living *E. coli*, provided a unique opportunity to explore the gain/loss of CK2^WT^ and CK2^K198R^ substrates at the site level. Specifically, we were interested in investigating whether our observed decreased preference for acidic residues at the +1 position in CK2^K198R^ resulted from the loss of existing *E. coli* substrates, the addition of completely new substrates, or a combination of the two. **Figure 3** shows a Venn diagram representing phosphorylation site overlap between CK2^WT^ and CK2^K198R^ indicating the latter; namely, approximately half of the CK2^WT^ sites were detected in the CK2^K198R^ mutant, yet over a third of the sites detected in the K198R mutant were not observed in the wild type kinase data set.

**Figure 3.**
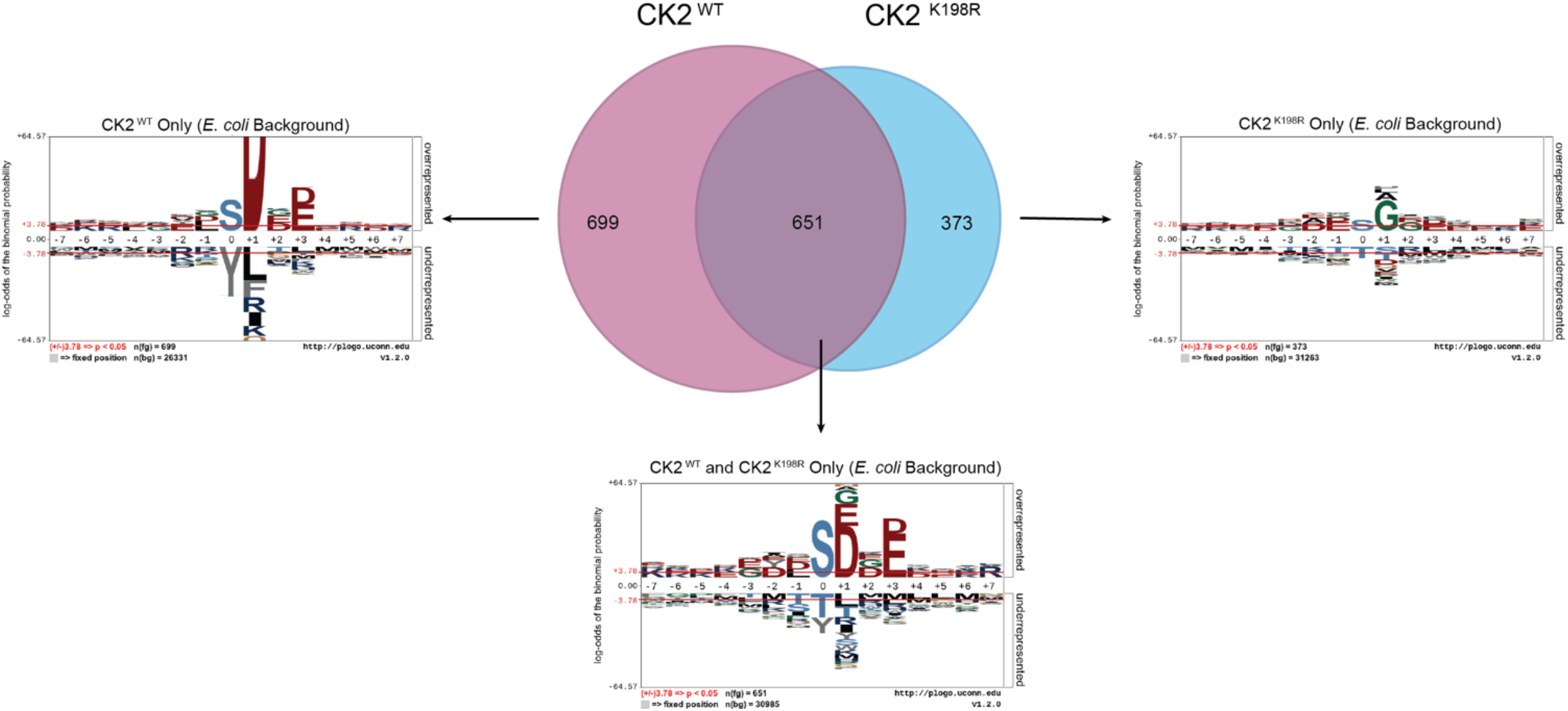
CK2^K198R^ phosphorylates an expanded complement of substrates. Motifs of phosphopeptides identified only in CK2^WT^, both CK2^WT^ and CK2^K198R^, or only in CK2^K198R^ ProPeL experiments. Phosphopeptides identified in both CK2^WT^ and CK2^K198R^ experiments tend to reflect the canonical CK2^WT^ motif. Those identified only in CK2^K198R^ experiments have a much wider complement of amino acids in the +1 position, indicating increased promiscuity in this position. The phosphoacceptor abundance is also altered, reflecting an increased preference for tyrosine in the K198R mutant.

To further explore motif differences between the three subsets noted in the Venn diagram, pLogos were generated for each subset (**Figure 3**). The pLogo generated from the set of phosphorylation sites unique to CK2^WT^ included a highly pronounced +1 aspartate specificity that became more attenuated in the overlapping data set and was entirely replaced by glycine (and to a lesser extent, alanine and leucine) in the sites unique to CK2^K198R^. The +3 acidic residues were similarly attenuated in the mutant kinase data set. Finally, with regard to the phosphoacceptor residue, the phosphorylation sites unique to CK2^WT^ showed a highly significant disfavoring of tyrosine residues relative to serine residues that was lost in the CK2^K198R^ mutant (which displayed a general neutrality to the phosphoacceptor residue).

Taken together these results point to a significant shift at the substrate level for the K198R mutant kinase, whereby the most canonical wild type CK2 phosphorylation sites are either lost or phosphorylated with decreased efficiency, coupled with a large increase in previously unphosphorylated sites.

### Predictions of CK2^WT^ and CK2^K198R^ phosphorylation sites

CK2^K198R^ is the most frequently occurring mutation in patients with OCNDS^4,15^. Substrate rewiring in this mutant, indicated by our specificity and *E. coli* substrate analyses, is likely to contribute to the etiology of OCNDS. In an effort to offer a potential mechanism for the role of CK2^K198R^ in OCNDS pathology we developed a predictive strategy for determining CK2^WT^ and CK2^K198R^ phosphorylation sites on human proteins based on our prior *scan-x* post-translational modification predictor^11^, but with the advantage of calculated probabilities of modification (see Methods). Due to the neurodevelopmental nature of OCNDS, in the present study we focused our predictions of potential CK2^WT^ and CK2^K198R^ phosphorylation sites to ion channels localized to the axon. The top ten predicted sites (from a list of over 3000 total sites) with the greatest change in probability between CK2^WT^ and CK2^K198R^ are listed in **Table 1** (a complete list of predicted sites is available in **Table S7**). Of the sites listed, two have an increased probability of modification by CK2^K198R^ and the remaining eight are less likely to be phosphorylated by CK2^K198R^. Additionally, four of the ten sites included in **Table 1** have been previously shown to be phosphorylated (with two additional sites being paralogs of phosphorylation sites). The two sites with the greatest increase in probability of modification by CK2^K198R^, S514 on the voltage-gated sodium channel 2A (SCN2A) and its paralog, S510 on the voltage-gated sodium channel 1A (SCN1A), are located in the I-II linker region. This region is modified by other kinases^16^, and increased phosphorylation of this segment is known to lead to a decrease in overall sodium channel activity.^17^ Another site of particular interest is S1126 on SCN2A, as this site falls in the ankyrin binding motif, and is a known CK2 phosphorylation site.^18–20^ Our predictions indicate that CK2^K198R^ is approximately 40% less likely to phosphorylate S1126 than CK2^WT^. Phosphorylation of S1126 by CK2 is essential for the accumulation of SCN2A in the axon through ankyrin^20^ and a loss of this accumulation could be relevant to the pathology of OCNDS patients possessing the CK2^K198R^ mutation.

## Discussion

The present study marks the first investigation into the impact of the OCNDS associated CK2^K198R^ mutation. Contrary to prior speculation, it is clear from our experiments that the CK2^K198R^ mutation does not simply cause a loss of function, or an inability to recognize substrates.^4,6^ We demonstrate that the K198R mutation results in a substantial shift in specificity by reducing the overall preference for acidic (D/E) residues at the +1 position, and by increasing preference for glycine residues at the same position.

While wild type CK2 has previously been shown to be capable of phosphorylating tyrosine residues^21–23^ the study also marks the first time a tyrosine phosphorylation motif has been identified for CK2, and, to our knowledge, the first time that a tyrosine phosphorylation motif has been identified for any kinase previously characterized as primarily phosphorylating serine and threonine residues. Interestingly, our study also revealed a highly significant shift in phosphoacceptor preference, with tyrosine residues being shifted from a disfavored state in CK2^WT^ to a neutral state in the CK2^K198R^ mutant, while threonine residues move in the precise opposite direction. Though the detection of a tyrosine phosphorylation motif for a kinase that disfavors tyrosine phosphoacceptors may appear to be a contradiction, it is important to note that the favoring and/or disfavoring of residues is calculated in the context of the background frequency of those residues. Thus, a phosphoacceptor may still experience significant phosphorylation, while being at a level far below what one would expect due to chance if all phosphoacceptors were phosphorylated at their expected background frequencies. Finally, our work highlights the importance of assessing phosphoacceptor specificity independently, as the shift in the specificity motif from CK2^WT^ to CK2^K198R^ differed between serine/threonine (both of which exhibited similar changes) and tyrosine, which added additional preferences for acidic residues upstream of the modification site in the mutant.

Among the advantages of our ProPeL methodology in assessing kinase specificity over alternate approaches is the fact that the method is carried out under physiological conditions in *E. coli*, thus allowing researchers to assess shifts in phosphorylation at the site level between kinases bearing different characteristics and/or mutations. Utilizing this strategy here we have shown that there exists overlap between CK2^WT^ to CK2^K198R^ at the substrate level, as well as considerable extension to new atypical CK2 substrates by the mutant kinase. The loss of highly canonical CK2 substrates coupled with the addition of new substrates provides a likely explanation for OCNDS etiology.

Another advantage of ProPeL in the present study is that the closed and finite system (i.e., living bacterial proteomes) allowed us to readily associate *scan-x* phosphorylation scores^11^ with actual probabilities of modification based on the observed modification state of all phosphorylatable residues in *E. coli*. Thus, for any score obtained for any peptide sequence, we could utilize the *E. coli* expressed proteome as a lookup table to correlate scores with probabilities. It is important to note that probabilities obtained utilizing this strategy represent an approximate lower bound on the probability as it is likely that many phosphorylatable sites in the *E. coli* lookup table are considered “negative” when they are indeed phosphorylated, while the converse is unlikely to be true due to our use of conservative peptide identification thresholds (i.e., “absence of proof is not proof of absence”).

Most kinases have multiple substrates throughout the cell, only a few of which will directly relate to a phenotype of interest. To investigate signaling changes that may underlie neurodevelopmental phenotypes associated with OCNDS, we focused on neuronal signaling in axonally localized ion channels, as CK2 has been shown to be highly enriched in the axon^20^ and is known to interact with axonal sodium and potassium channels.^24,25^ Further, these ion channels are found to be frequently mutated in both epilepsies and neurodevelopmental disorders, with one recent study reporting 5% of 8,565 participants carrying a mutation in axonally localized voltage gated sodium channels.^26,27^ Of 3000+ sites predicted be differentially phosphorylated by CK2^WT^ and CK2^K198R^, the ten sites with the greatest differences present several interesting candidates for hypothesis generation. The two sites with the greatest increased probability of modification by CK2^K198R^, S514 on SCN2A and S510 on SCN1A, are located in the I-II linker, a region known to be modified by other kinases including PKA and PKC.^16^ Interestingly, increased phosphorylation of sites in this region reduces the overall activity of the channel^17^ potentially indicating that the CK2^K198R^ may decrease activity of SCN1A and SCN2A through increased phosphorylation of the I-II linker region. Additionally, our predicted 4^th^ most differentially modified site was S1126 on the voltage-gated sodium channel (SCN2A), a site that is not only already known to be phosphorylated by CK2, but also bears great importance to the proper accumulation of SCN2A to the axon via ankyrin binding. Specifically, ankyrin-mediated binding links SCN2A to the cytoskeleton as well as other scaffolding proteins to localize the channel to the proper location in the plasma membrane.^20^ Without this precise accumulation, it has been shown that propagation of action potentials is significantly impaired.^28^ Furthermore, this prediction is especially pertinent to the study of OCNDS as SCN2A is the predominant isoform of sodium channel found in neurons early in development^29^, and as such any defect in accumulating this ion channel could be detrimental for the proper formation of neural circuits. Additional investigation of the interaction between CK2^K198R^ and voltage gated sodium channels is clearly warranted. It is our intention to eventually make publicly available a global proteomic CK2^K198R^ versus CK2^WT^ phosphorylation predictor, as the underlying mechanism by which OCNDS leads to disease is almost certain to be multifactorial given the phenotype is not limited to the central nervous system.^1,4,15^

Finally, as our group and others previously demonstrated that the Cushing’s Syndrome mutation PKA^L205R^ results in altered substrate specificity^30–32^, the present study provides another data point to suggest that there exists a subset of disease-linked kinase mutations whose deleterious effects can be traced to subtle shifts in substrate specificity rather than broad alterations of kinase activity. The genomic revolution is likely to reveal that we are only at the tip of the proverbial iceberg, as patients with novel mutations are more frequently being identified. By coupling simple experimental strategies for kinase specificity determination with computational approaches to predict substrates, our hope is to generate testable hypotheses regarding underlying mechanisms of disease that can lead us to potential therapeutics faster than ever before.

## Acknowledgements

This work was funded in part by a grant awarded to DS by the National Institute of Neurological Disorders and Stroke (1R21NS096516).

## Author Contributions

AVT and DS conceived of the study. DMC and DS designed the ProPeL experiments. DS designed the computational scoring methodology. DMC and JCL performed the experiments. NQP and JPO, implemented the computational scoring methods. DMC analyzed the data. AVT, JLB, JCL, and DS contributed materials, resources, and/or analysis tools. DMC and DS wrote the manuscript. All authors helped edit the final manuscript.

## References

1. Okur V, Cho MT, Henderson L, et al. De novo mutations in CSNK2A1 are associated with neurodevelopmental abnormalities and dysmorphic features. Hum. Genet. 2016;135(7):699–705.

2. Pinna LA. Casein kinase 2: An ‘eminence grise’ in cellular regulation? Biochim. Biophys. Acta BBA - Mol. Cell Res. 1990;1054(3):267–284.

3. Jiménez JS, Benítez MJ, Lechuga CG, et al. Casein kinase 2 inactivation by Mg2+, Mn2+ and Co2+ ions. Mol. Cell. Biochem. 1995;152(1):1–6.

4. Chiu ATG, Pei SLC, Mak CCY, et al. Okur-Chung neurodevelopmental syndrome: Eight additional cases with implications on phenotype and genotype expansion. Clin. Genet. 2018;93(4):880–890.

5. Nolen B, Taylor S, Ghosh G. Regulation of Protein Kinases: Controlling Activity through Activation Segment Conformation. Mol. Cell 2004;15(5):661–675.

6. Sarno S, Vaglio P, Meggio F, et al. Protein Kinase CK2 Mutants Defective in Substrate Recognition PURIFICATION AND KINETIC ANALYSIS. J. Biol. Chem. 1996;271(18):10595–10601.

7. Lubner JM, Church GM, Chou MF, Schwartz D. Reprogramming protein kinase substrate specificity through synthetic mutations. bioRxiv 2016;091892.

8. Lubner JM, Balsbaugh JL, Church GM, et al. Characterizing Protein Kinase Substrate Specificity Using the Proteomic Peptide Library (ProPeL) Approach. Curr. Protoc. Chem. Biol. 2018;10(2):e38.

9. Cox J, Mann M. MaxQuant enables high peptide identification rates, individualized p.p.b.-range mass accuracies and proteome-wide protein quantification. Nat. Biotechnol. 2008;26(12):1367–1372.

10. Chou MF, Prisic S, Lubner JM, et al. Using Bacteria to Determine Protein Kinase Specificity and Predict Target Substrates. PLOS ONE 2012;7(12):e52747.

11. Schwartz D, Chou MF, Church GM. Predicting Protein Post-translational Modifications Using Meta-analysis of Proteome Scale Data Sets. Mol. Cell. Proteomics 2009;8(2):365– 379.

12. Macek B, Gnad F, Soufi B, et al. Phosphoproteome Analysis of E. coli Reveals Evolutionary Conservation of Bacterial Ser/Thr/Tyr Phosphorylation. Mol. Cell. Proteomics 2008;7(2):299–307.

13. Soares NC, Spät P, Krug K, Macek B. Global dynamics of the Escherichia coli proteome and phosphoproteome during growth in minimal medium. J. Proteome Res. 2013;12(6):2611–2621.

14. O’Shea JP, Chou MF, Quader SA, et al. pLogo: a probabilistic approach to visualizing sequence motifs. Nat. Methods 2013;10(12):1211–1212.

15. Owen CI, Bowden R, Parker MJ, et al. Extending the phenotype associated with the *CSNK2A1‐* related Okur–Chung syndrome—A clinical study of 11 individuals. Am. J. Med. Genet. A. 2018;176(5):1108–1114.

16. Berendt FJ, Park K-S, Trimmer JS. Multisite Phosphorylation of Voltage-Gated Sodium Channel α Subunits from Rat Brain. J. Proteome Res. 2010;9(4):1976–1984.

17. Cantrell AR, Catterall WA. Neuromodulation of Na+ channels: an unexpected form of cellular plasticity. Nat. Rev. Neurosci. 2001;2(6):397–407.

18. Garrido JJ, Giraud P, Carlier E, et al. A targeting motif involved in sodium channel clustering at the axonal initial segment. Science 2003;300(5628):2091–2094.

19. Lemaillet G, Walker B, Lambert S. Identification of a conserved ankyrin-binding motif in the family of sodium channel alpha subunits. J. Biol. Chem. 2003;278(30):27333–27339.

20. Bréchet A, Fache M-P, Brachet A, et al. Protein kinase CK2 contributes to the organization of sodium channels in axonal membranes by regulating their interactions with ankyrin G. J. Cell Biol. 2008;183(6):1101–1114.

21. Vilk G, Weber JE, Turowec JP, et al. Protein kinase CK2 catalyzes tyrosine phosphorylation in mammalian cells. Cell. Signal. 2008;20(11):1942–1951.

22. Basnet H, Bessie Su X, Tan Y, et al. Tyrosine phosphorylation of histone H2A by CK2 regulates transcriptional elongation. Nature 2014;516(7530):267–271.

23. Marin O, Meggio F, Sarno S, et al. Tyrosine Versus Serine/Threonine Phosphorylation by Protein Kinase Casein Kinase-2 A STUDY WITH PEPTIDE SUBSTRATES DERIVED FROM IMMUNOPHILIN Fpr3. J. Biol. Chem. 1999;274(41):29260–29265.

24. Hien YE, Montersino A, Castets F, et al. CK2 accumulation at the axon initial segment depends on sodium channel Nav1. FEBS Lett. 2014;588(18):3403–3408.

25. Kang S, Xu M, Cooper EC, Hoshi N. Channel-anchored protein kinase CK2 and protein phosphatase 1 reciprocally regulate KCNQ2-containing M-channels via phosphorylation of calmodulin. J. Biol. Chem. 2014;289(16):11536–11544.

26. Lindy AS, Stosser MB, Butler E, et al. Diagnostic outcomes for genetic testing of 70 genes in 8565 patients with epilepsy and neurodevelopmental disorders. Epilepsia 2018;59(5):1062–1071.

27. Brunklaus A, Lal D. Sodium channel epilepsies and neurodevelopmental disorders: from disease mechanisms to clinical application. Dev. Med. Child Neurol. 2020;62(7):784–792.

28. Spratt PW,Ben-Shalom R, Keeshen CM, et al. The autism-associated gene Scn2a contributes to dendritic excitability and synaptic function in prefrontal cortex. Neuron 2019;103(4):673-685.e5.

29. Kaplan MR, Cho MH, Ullian EM, et al. Differential control of clustering of the sodium channels Na(v)1.2 and Na(v)1.6 at developing CNS nodes of Ranvier. Neuron 2001;30(1):105–119.

30. Lubner JM,Dodge-Kafka KL, Carlson CR, et al. Cushing’s Syndrome mutant PKAL205R exhibits altered substrate specificity. FEBS Lett. 2017;591(3):459–467.

31. Walker C, Wang Y, Olivieri C, et al. Cushing’s syndrome driver mutation disrupts protein kinase A allosteric network, altering both regulation and substrate specificity. Sci. Adv. 2019;5(8):eaaw9298.

32. Bathon K, Weigand I, Vanselow JT, et al. Alterations in Protein Kinase A Substrate Specificity as a Potential Cause of Cushing Syndrome. Endocrinology 2019;160(2):447–459.

